# Integrating Electrical Components into a Printed Self-folding Cuff Electrode for Chronic Peripheral Nerve Interfaces

**DOI:** 10.64898/2026.03.16.712029

**Authors:** Lukas Hiendlmeier, Defne Tüzün, Hannah Tillert, Anton Dalichau, Mehmet Öztürk, Yannick Günzel, Hannes Kübler, Francisco Zurita, George Al Boustani, Jose Zariffa, Einat Couzin-Fuchs, George G. Malliaras, Amparo Güemes, Bernhard Wolfrum

## Abstract

Neuroelectronic thin-film implants hold promise for advancing fundamental understanding of the peripheral nervous system and offer potential for targeted treatments using specific stimulation. However, challenges in establishing robust and durable connections to soft and flexible implants limit their widespread adoption and long-term utility. Here, we present a novel method for integrating rigid electrical components, such as a standard USB-C connector, directly into a printed stretchable self-folding cuff electrode for chronic peripheral nerve interfacing. Our multi-material printing approach provides a gradual stiffness transition, effectively mitigating common failure points associated with mechanical stress at soft-rigid boundary. We demonstrate the integration of a wireless stimulation circuit and a robust USB-C implantable port, offering a ‘plug-and-play’ solution for stable chronic electrophysiology experiments. Chronic implant studies in free-running locusts with USB-C connectors show reliable nerve recordings, capturing behavioral differences. The concept is further validated as a transcutaneous implanted port in rats for vagus nerve recordings. This work addresses a critical bottleneck in neurotechnology by enabling robust connectivity for implanted devices, which is essential for advancing peripheral nervous electrophysiology experiments in freely moving small vertebrates and insects.

## Introduction

Neuroelectronic implants, that interface with neural tissue in the central and peripheral nervous systems (PNS), offer promising targeted, long-term treatments.^1^ While clinically applied for conditions like epilepsy (vagus nerve), sleep apnoea (hypoglossal nerve), or bladder control (sacral root),^2–4^ current PNS implants often employ non-specific electrical stimulation of large nerves via distant, big electrodes. These non-specific interfaces contrast with the nervous system’s precise organ innervation, revealing important opportunities for targeted neural interfacing and modulation of still poorly understood pathways (e.g., vagal nerve effects on glucose regulation and inflammation or the innervation of brown adipose tissue).^5–7^ Developing new neuroelectronic implants for stable, localised recording along smaller nerves with minimal tissue damage is thus crucial. Such advancements would not only enhance fundamental understanding of PNS organ influence and aid diagnostics, but also lead to substantial medical breakthroughs.^8,9^

Despite the need for improved neuroelectronic devices, the development of new implants still requires advancement on the following key challenges:^10,11^

1. Reducing tissue damage and foreign body response (e.g., optimising device mechanics and handling).^12–15^
2. Enhancing the electrode-electrolyte interface for low impedance and high charge injection capacity of microelectrodes.^15–18^
3. Improving device stability by addressing the corrosion of the electrode sites or delamination of either the conductive layer or the passivation layer.^19–21^
4. Addressing connections and setup failures such as unreliable contacts or broken conductor traces.^22,23^

While we predominantly address the fourth challenge within this work, many recently developed implants focus on the first one. To enable closer proximity to the targeted neuronal tissue for higher spatial selectivity and signal quality, the tissue damage caused by the implantation and the foreign body response must be minimised. Therefore, many developed implants are made from thin-film polymers that are conformal, stretchable, and soft, even incorporating hydrogels, to reduce mechanical forces applied to the tissue.^18,24–36^ For example, we previously developed stretchable and self-folding cuff electrodes, that allow easy handling and implantation in acute experiments.^37^ However, their reliance on zero insertion force (ZIF) connectors, often employed in flexible electronics and implants,^34,38^ can become problematic for chronic applications due to mechanical damage of the metal layer, leading to connection failures (fourth challenge). This highlights a general issue with the connection between soft and flexible implants and rigid electronics.^26,39–41^ It is often caused by the mechanical transition from flexible or stretchable materials to rigid electronic and electromechanical components.^42,43^

Typical connection methods for neural devices to the rigid electrical circuit include: conductive glueing,^44^ wire bonding,^45^ low-temperature soldering,^46^ and anisotropic conductive film or paste bonding.^29,47^ These methods often involve complex multi-material assemblies and present trade-offs between electrical performance and mechanical stability, introducing potential failure points. Even commonly used thick silicone encapsulation, which aims to reduce local stress and protect material interfaces from moisture-related corrosion and delamination, mainly mitigates risks rather than completely solving them.^44,46,48^

Beyond the soft-rigid interface, transmitting power and signals for chronic experiments presents another challenge.^39^ While fully implanted wireless systems offer benefits and are usually used for clinical settings, they are often impractical for research involving neuronal recording from small animals, since they are typically too large, too costly, or lack sufficient bandwidth.^49,50^ This leads to the reliance on transcutaneous connections. However, these external connections carry risk of infection and need to be mechanically anchored. This necessitates robust and reliable transcutaneous ports that minimise open wounds and transfer the forces from cables away from the implant into the surrounding tissue (skin or bones). ^22,51^

This research presents a method to overcome soft neuroelectronic device connectivity issues through integrated connection ports, thus enabling reliable chronic experiments. Our example demonstrates embedding a standard USB-C connector directly into printed stretchable self-folding electrodes for chronic recording. This approach aims to minimise the number of assembly steps and material variations. A controlled gradual stiffness transition from soft to rigid regions makes the interface mechanically robust. Additionally, the integrated connector enables reliable plugging and unplugging with multi-channel devices, while allowing the fixation of the port on or through the animal’s skin.

## Results

### Fabrication process

The fabrication process for the self-folding electrodes, as well as the one with integrated electronic components, is shown in Figure 1. To fabricate the bare self-foldable electrodes (Figure 1a), frontal photopolymerization (FPP) with UV-curable acrylated printing resins is used to produce the electrode substrate. The FPP process involves exposing a pattern through a transparent glass slide to cure the liquid resin, allowing for the definition of a 2D structure. The exposure time determines the thickness of the polymerised layer, influencing the bending stiffness of different structural parts, such as folding regions. Since this exposure does not completely cure the uppermost layer, additional printing resins, such as a superabsorbent polymer (SAP) hydrogel, can be printed on top. The resins create strong covalent bonds to the previously printed layer, as they are radical-polymerising acrylates, and not all bonds are initially activated during the printing.^52,53^ Thereby, we successfully fabricate the electrode substrate with a defined 2.5D structure that incorporate functionalities, such as varying thickness, bending stiffness, and self-folding upon contact with water. After detaching from the substrate, the structure is flipped, and a thin layer of gold is deposited onto the flat surface, forming a µ-cracked, stretchable conductive layer. The conductive film in the areas between the conductive traces is removed via laser ablation, and a passivation layer of the same printing resin is spin-coated. After the passivation layer is cured, the final devices are cut out.

**Figure 1:**
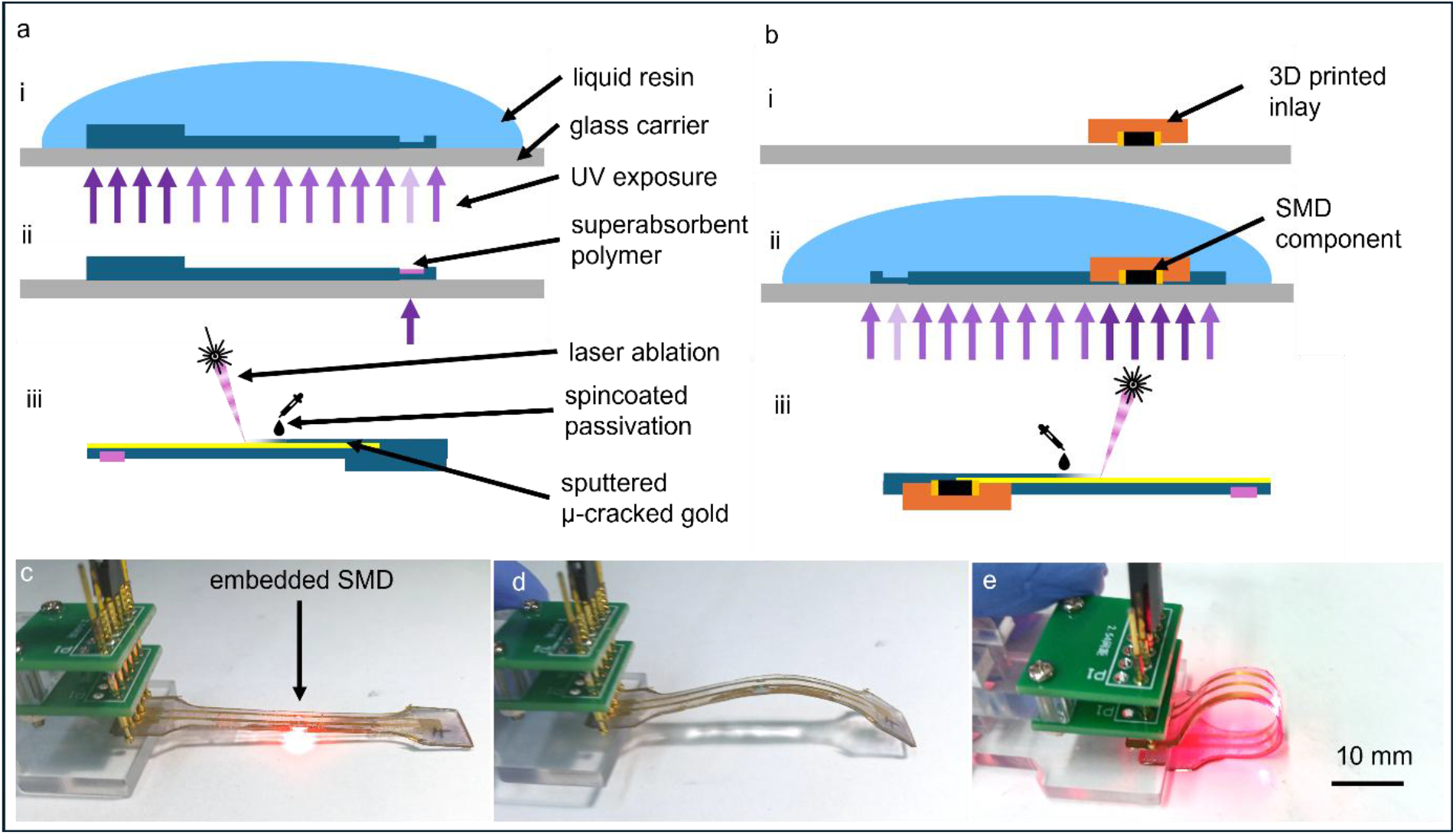
Fabrication schematics of the self-folding cuff electrodes with and without embedded electrical components. (a) Schematic illustrating the fabrication workflow of the electrode without electrical components. (i) The electrode substrate is printed by FPP from liquid flex-resin layers with varying thicknesses. (ii) A superabsorbent polymer layer facilitating self-folding is polymerised on top. (iii) The electrode is formed by gold sputtering and laser ablation, and the feed lines are isolated using flex-resin. (b) Integration of electronic components (e.g. SMDs). (i) A 3D-printed inlay positions the SMD, (ii) which is then embedded by curing with flex-resin. (iii) Sputtered gold connects the contact pads of the SMD components after the removal of the glass carrier. (c) Example of a functional test structure with an embedded resistor and LED. (d) Illustration of connection failure when the stiffness of the 3D-printed inlay is the same as the flexible layer, causing cracking under strain. (e) Demonstration of successful integration with an inlay stiffness optimised to mitigate cracking and maintain functionality.

We adjusted the fabrication process to integrate rigid electromechanical components into the self-foldable cuff electrodes, as shown in Figure 1b. Since the substrate is flipped after printing, we incorporate electronic components into the structure before connecting those components with the laser-patterned gold film. An inlay is 3D printed using stereolithography (SLA) and features voids designed to hold and align surface-mount device (SMD) components. These components are placed in the voids such that all exposed contact pads are aligned in a single plane. The inlay is then placed on a glass slide carrier with the contacts facing down, filled with liquid resin, and exposed using FPP, following the same fabrication method as the self-folding electrodes. Since the previously printed inlay is not fully cured, it cures simultaneously with the new layer, forming a continuous material akin to the layer-to-layer adhesion process observed in SLA printing. The complete print, including the components, is then lifted off the glass carrier and flipped around, exposing the contact pads on top. These pads can be manually cleaned of any remaining resin that may flow under the parts during the process. Alternatively, the inlay is placed on a thin layer of polyvinyl alcohol (PVA) glue while positioned onto the glass carrier to prevent resin leakage, which can be washed away with water. The sputtered gold successfully connects the components without damaging the electronic parts during the laser patterning of the circuit. To validate electrical connectivity and structural integration, we fabricated a simple circuit containing a resistor and a light-emitting diode (LED), with the connection traces formed by the gold thin film embedded in the printed structure. As shown in Figure 1c, the resistor and LED were successfully integrated into the circuit and illuminated.

### Mechanical characterisation of integration

When a sample made from flexible printing resin with mounted components and the inlay being printed from the same flexible resin is bent with a radius of approximately 5 mm, the circuit breaks and undergoes irreversible electrical failure (Figure 1d, Video SI1). This failure is likely due to tensile loads acting on the connections during bending.

To investigate this, we performed tensile tests on different types of samples: blank samples consisting of gold feedlines embedded in flexible resin, and samples with two 0-Ω resistors embedded between the feedlines using a 3D printed inlay (Figure 2a). Along the stress-strain curves, the resistance of the samples was measured as they were cyclically stretched. Figure 2b shows the stress-strain relationship of the tested samples over the first, second, and tenth cycles, stretched by 20% strain relative to the initial gauge length. The material exhibits pronounced hysteresis, a characteristic of the flexible resin used in this application. The measured E_M_ of the blank sample (blue) was 3.3 ± 0.3 (n=33) MPa.

**Figure 2:**
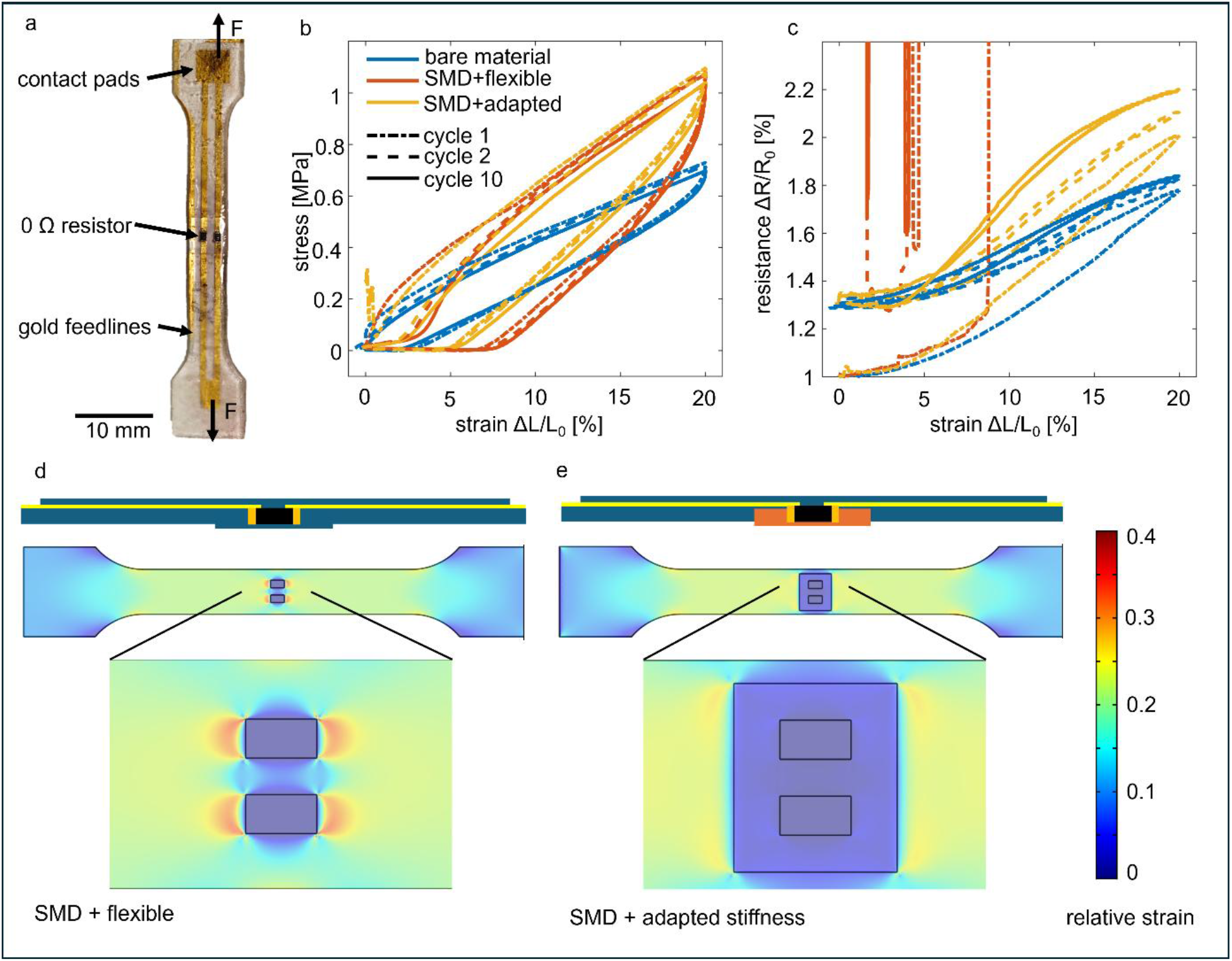
Electromechanical characterisation of the integrated component. (a) Picture of a test sample (b) The stress-strain relation during cyclic tensile tests of the samples: blank sample of the bare material without the inserted component (blue), SMD resistor embedded in flexible resin (orange), and SMD resistor with the stiffness-adapted insert (yellow). The first, second, and tenth cycles are shown with different line styles. (c) The relative change in resistance of the gold track with two 0 Ω resistors during cyclic tensile tests. (d) A FEM simulation displaying the relative local strain in the direction of the tension with flexible inlay. (e) A FEM simulation as (d) with a stiffness-adapted inlay.

Figure 2c illustrates the relative change in resistance compared to the initial state during stretching cycles. The blue curve represents the blank sample, which shows an expected increase in resistance while being stretched due to the µ-cracked gold film. After the first cycle, the resistance did not return to its original value but increased slightly because of the formation of the µ-cracks. The resistance stabilised during the subsequent cycles. The orange curve represents a sample that includes the SMD components with an inlay made from the same flexible material as the rest of the device. This configuration exhibited an irreversible increase in resistance after strain of approximately 10% was applied to the material. When stretched, the gold connections crack and do not reliably recover to the initial resistance value upon release.

To identify the failure location, we conducted a finite element method (FEM) simulation (Figure 2d). The relative strain tensor of the gold layer in the load direction reveals high strain at the transition between the rigid SMD component and the surrounding flexible material. This, combined with poor mechanical adhesion due to differing materials, presumably leads to electrical connection failure at this interface between the component and the resin when applying strain.^43^

To address this issue and distribute strain more evenly across the weak interface, we printed the inlay around the electrical component using a stiffer printing resin Medical Print Clear (E_M_ ∼ 400 MPa).^54^ This modification creates a more gradual transition between the rigid SMD component (E_M_ ∼ GPa) and the stretchable flexible resin (E_M_ ∼ 3 MPa). Similar principles have been documented in the literature.^55,56^ As shown in Figure 2e, the increased strain is directed away from the interface of the SMD component towards the interface between the two resins. Since both resins are acrylates, they cure together, forming a strong adhesion that is less susceptible to cracking. The tensile tests of these modified samples demonstrate the circuit remains conductive (yellow curve, Figure 2c) throughout the cycles. Notably, the adapted sample shows a higher resistance increase compared to the blank sample during stretching (Figure 2c). This increase can be attributed to the introduction of the stiff insert, which effectively reduces the length of the flexible material being stretched. Additional test samples are presented in the Supporting Information (SI2). As Figure 1e and Video SI3 illustrate, the LED test circuit with adapted stiffness remains functional even when bent with a radius of approximately 5 mm.

### Wireless stimulator

After successfully showing the integration of SMD components into the fabrication process, we integrated a near-field communication (NFC) circuit into the printed self-folding cuff electrode design for neuronal stimulation. Figure 3a illustrates the schematic circuit around the NFC NTAG 5 chip, which includes an SMD antenna and additional capacitors for power harvesting. The antenna circuit is tuned to the NFC frequency of 13.56 MHz using a capacitor, as in conventional PCBs (Figure 3b). The output voltage of the printed circuit, a train of 200 µs pulses with a period of 10 ms, is shown in Figure 3c. This output is then connected to self-folding cuff electrodes via gold tracks, displayed in Figure 3d. This device mounts on a locust’s back; its cuffs interface with the left and right N5 nerves, controlling the hind legs. As shown in Figure 3e and Video SI4, the nerve can be stimulated via the implanted device using the NFC field, eliciting a leg movement. The range of the NFC field is enhanced by focusing it through a matched resonance coil placed on top of it. The demonstration on the locust, does not resolve the typical application where power and data should be transmitted wirelessly through the skin. Therefore, we printed the stretchable circuit with integrated LEDs as optical indicators for the transmission, as shown in Figure 3f. Figure 3g illustrates the implanted LED circuit in a chicken leg, where transcutaneous energy transfer and communication can be achieved through the skin (Video SI5).

**Figure 3:**
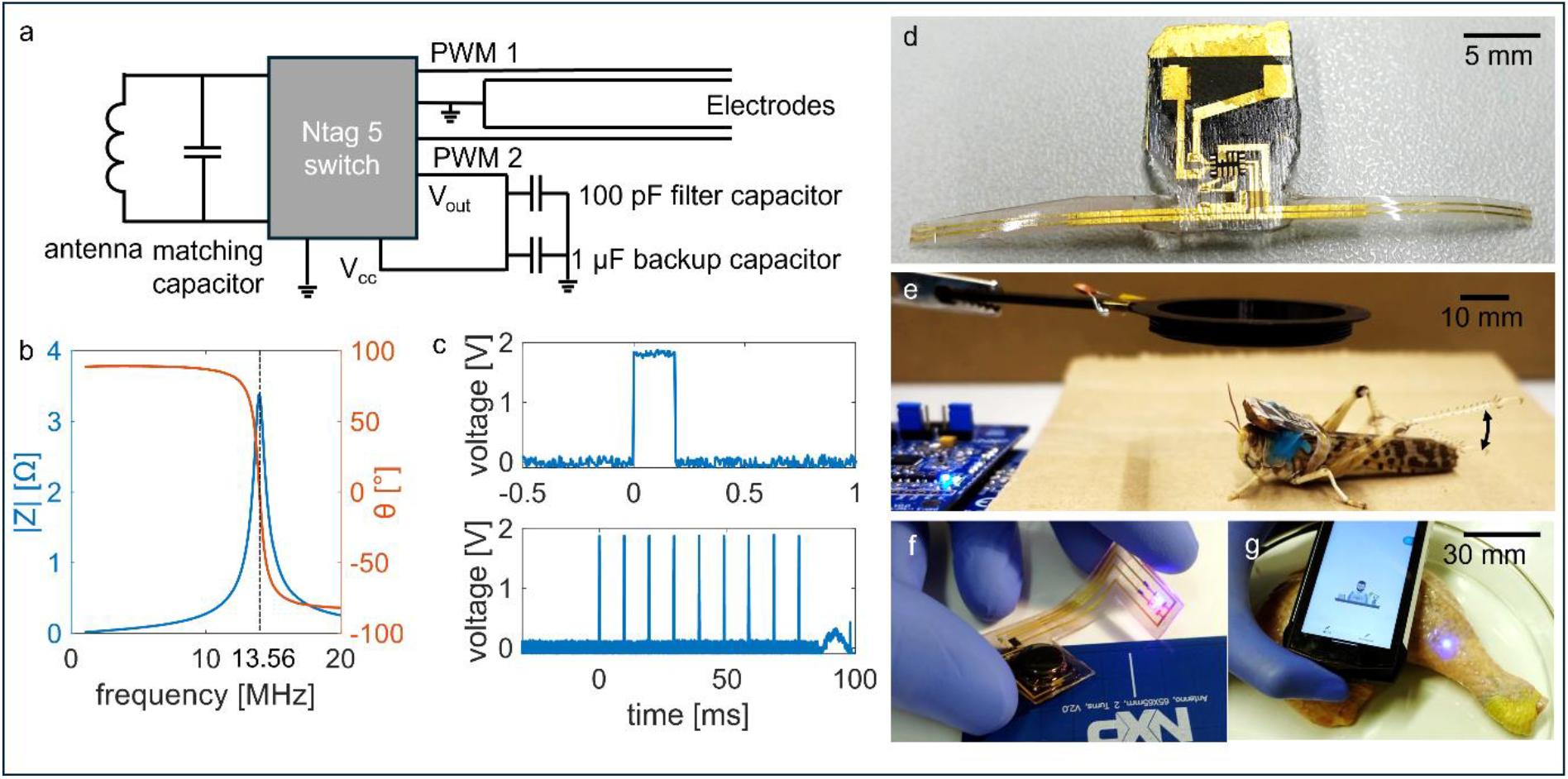
Integrated self-folding electrodes with an NFC stimulator. (a) Circuit schematic of the self-folding electrodes integrated with an NTAG 5 chip, an SMD antenna, and additional capacitors to allow power harvesting. (b) Resonance of the integrated NFC antenna matched to 13.56 MHz. (c) Generated voltage pulses for stimulation. (d) Image of the self-folding electrodes with the fully-integrated NFC stimulator circuit. (e) The device is implanted to interface with the hind legs of the locust. (f) Fully integrated flexible NFC circuit driving an LED. (g) Device shown in f, implanted into a chicken leg, powered by a mobile phone.

### Embedding a connector

Unlike the demonstrated neuronal stimulation, where wireless systems are readily implemented, multichannel neuronal recording presents significant challenges. These difficulties arise from high power consumption, limited bandwidth, and the resulting high cost and weight of the electronics, rendering them unsuitable for research in small animals such as rodents. Therefore, cables remain the standard for neural recordings in small animal research, facilitating the transition of power and data transmission to/from external electrophysiology equipment. For that, ideally, stable transcutaneous connectors must be integrated into the implants. To achieve this, we utilised commercially available USB-C connectors (Figure 4a) for integration into the printed cuff electrode as implantable transcutaneous connectors. USB-C connectors were selected because of their widespread availability, lightweight design, capability to support up to 24 channels, integrated housing for grounding, and extensive durability for millions of mating cycles.

**Figure 4:**
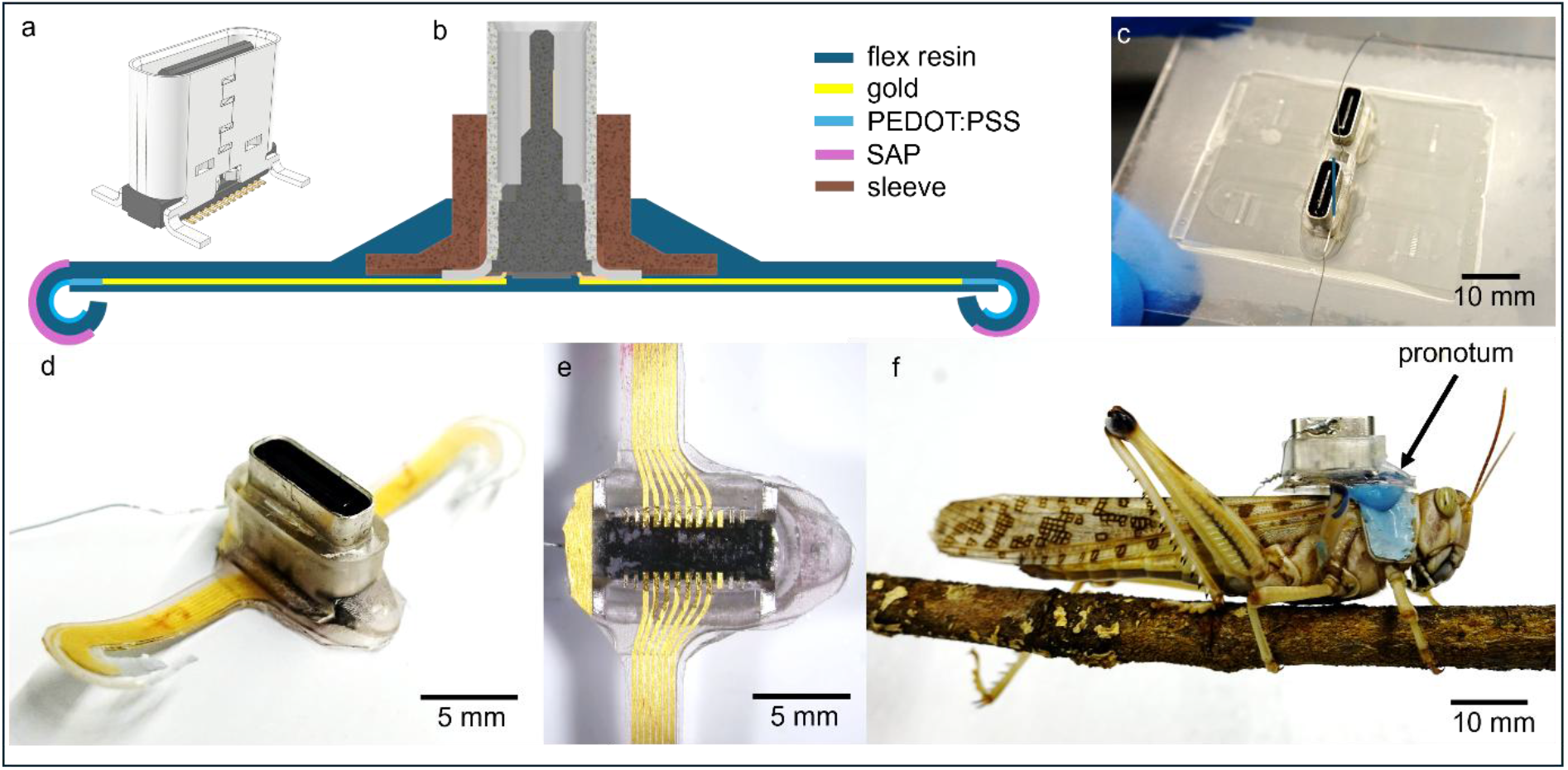
Integration of self-folding electrodes with a USB-C connector for implantation. (a) Schematic of a commercially available USB-C connector featuring 24 pins. (b) Cross-sectional schematic of the self-folding electrodes integrated with a USB-C connector. Colours indicate material composition: brown, stiffer resin; dark blue, flexible resin; yellow, gold feed lines; light blue, PEDOT:PSS electrodes; pink, superabsorbent hydrogel. (c) Image of 3D-printed resin electrode substrates with integrated USB-C connectors before peeling them off. (d) Final device with an integrated stainless-steel ground/reference electrode. (e) Bottom view of the final device showing gold feed lines connecting the electrodes to the contact pads of the embedded USB-C connector. (f) Cybernetic insect with an implanted device interfacing both N5 nerves of the hindlegs of a locust, with the connector mounted on the pronotum using UV-curable glue.

The connector is integrated into a separately 3D-printed sleeve made from the stiff resin, which is then positioned on a glass slide, allowing the flexible resin to cure around it and form the self-folding cuff substrate. The sleeve stabilises the connector within the printed structure and serves as a stiffness transition between the rigid connector and the flexible feed lines. A schematic cross-section of the connector in the sleeve integrated with self-folding cuff electrodes is shown in Figure 4b.

Figure 4c presents the printed self-folding substrate with integrated USB-C connectors before it is lifted off the glass carrier. The final device, featuring two separate cuff electrodes for the locust’s left and right hind legs, is depicted in Figure 4d. Figure 4e shows the gold film tracks connecting to the pads of the USB-C connector. The electrode tips are produced by spin-coating poly(3,4-ethylenedioxythiophene) polystyrene sulfonate (PEDOT:PSS) over the gold layer and on the tips where no gold was deposited. This conductive polymer is patterned during the same laser micromachining step as the gold layer. PEDOT:PSS substantially reduces the impedance compared to gold, therefore enhancing recording quality (impedance spectra of PEDOT:PSS vs gold electrodes are shown in Figure SI6).^57^

This built-in connector can be attached to the locust’s pronotum (the dorsal side of the prothorax) using UV-curable dental glue (blue), as shown in Figure 4f. This design transfers the stress from the nerve interface at the electrode entry points when plugging and unplugging the wired amplifier. Video SI7 shows a freely walking locust with the amplifier connected via a flexible PCB cable. Weighing approximately 0.8 g without the amplifier, the device is lighter than a typical adult desert locust (2 g). At this weight, the implant is not expected to affect their behaviour, since the females are used to carrying the males on their backs.

### Chronic nerve recordings

To evaluate the long-term functionality of the self-folding cuff electrodes integrated with a USB-C connector, we performed chronic recordings of nerve activity. The cuffs were implanted bilaterally onto the N5 nerves innervating the hind legs with successful recordings of both legs simultaneously lasting for several weeks from six locusts. Four additional animals were excluded from the analysis because of self-inflicted implant removal or infections that occurred within the first three days after surgery. Each implant featured up to ten recording channels, five on each cuff, arranged as ring electrodes in a linear configuration along the cuff with 150 µm spacing. The recordings were made in a monopolar configuration against a stainless-steel ground electrode placed in the animal’s abdomen. All recorded data was bandpass filtered between 200 and 4000 Hz unless stated otherwise. To analyse and display the data, we calculated the median value of the signals within each cuff and the bipolar signal between the outermost channels of each cuff.

Figure 5a illustrates the number of functional channels, defined as having an impedance < 1 MΩ. It should be noted that one of the potential ten channels consistently exhibited intermittent impedance from the outset across multiple animals because of an unstable contact within the connector cable, effectively reducing the reliable channel count to nine. In some devices, an additional electrode channel gradually became non-functional over time. While a clear failure mode for this breakage was not identified, potential causes include instability within a connector channel or damage to the gold conductor tracks leading to the connector, both likely attributed to mechanical stress. Only locust 3 (yellow) shows a steep decline in the number of functional channels, which occurred after day 7, all of which are located on the same side of the animal. This decline resulted from mechanical damage to the electrode feedlines, visually confirmed through further inspection, which was likely caused by the locust trying to rip it off using its legs. The remaining channels on the contralateral side continued recording reliably. Figure 5b displays the mean impedance of the functional channels for each animal. A moderate increase in impedance was observed during the first week post-implantation, after which values generally stabilised. This trend is consistent with tissue responses such as encapsulation or regrowth around the electrodes, and it indicates stable interfacing following the initial post-surgical period.

**Figure 5:**
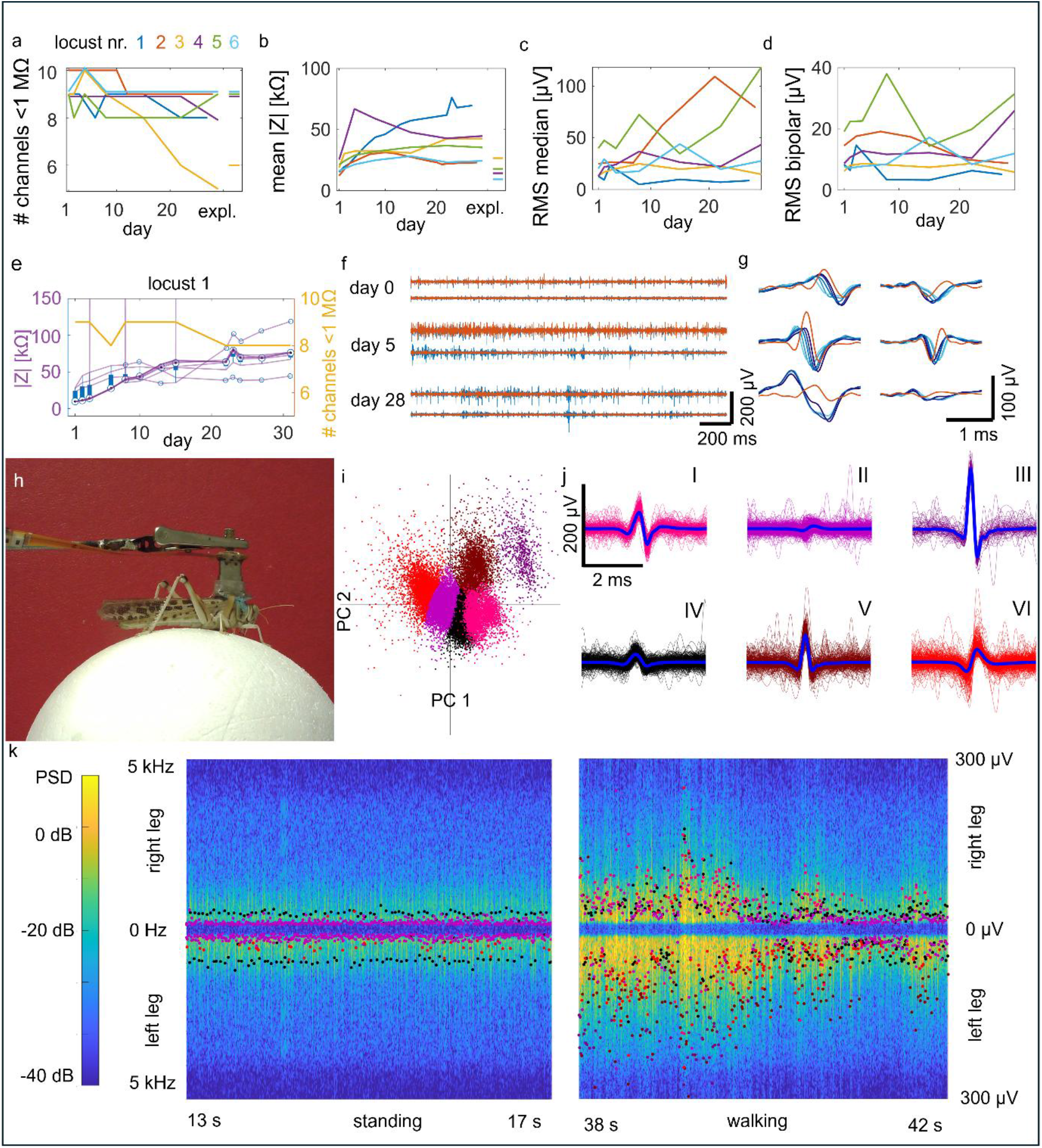
Chronic neural recording capability of self-folding electrodes implanted in locusts on both nerve 5 (N5) innervating the hind legs. (a-d) Data from six locusts chronically implanted for up to 30 days. (a) Number of functional channels with impedance below 1 MΩ at 1kHz. The final data point shows the number of working channels measured after explantation on day 30. (b) Mean impedance of all functional channels per animal. (c) RMS of the median signals recorded from one cuff of a 4-second trace of movement. (d) RMS of the calculated bipolar signal. (e-g) representative signal traces of one locust over time. (e) Number of working channels and impedances of electrode channels over 30 days post-implantation in locust 2. (f) Representative neural signal traces from both nerves, showing the median signal (blue) for each cuff and the calculated bipolar signal (orange) between the furthest apart channels in the cuff. (g) Close-up of neural signals (e.g., CAP) shown in (f). The blue traces represent signals from individual channels, with colour gradient matching their order within the cuff, and the orange trace shows the calculated bipolar signal. (h) Image of the recording setup, allowing neural data to be acquired from moving animals placed on an air-elevated polystyrene foam ball. The implanted device is connected to the acquisition system via a flexible cable, here in orange. (i) PCA of the bipolar signals with identified spikes from the median trace, revealing distinct clusters. (i) Bipolar waveforms from clusters identified in (h), differing in shape and amplitude. (k) Spectrograms of neural activity from left and right legs during standing and walking. Overlaid dots indicate identified spikes, colours denote the identified clusters in (i), and their height denotes the amplitude of the spike derived from median data.

Following the analysis of the electrode functionality and impedance trends over time, we next assessed the recorded signals. Figures 5c and 5d show the root-mean-square (RMS) values of the recorded signals from cuff electrodes implanted in six different locusts. The RMS values are extracted from a 4-second movement period, identified from video footage. Figure 5c displays the RMS of the median signal; in contrast, Figure 5d shows the RMS of the bipolar signal. As expected, median RMS values were generally higher than bipolar RMS values. This difference is caused by the median RMS’s sensitivity to non-neural signals, such as electromyography (EMG) signals generated during movement. In contrast, bipolar RMS suppresses common-mode signals, providing a more conservative estimate of neural recordings. Across animals, trends in RMS amplitude varied. In locust 1, both RMS values declined after day 3, aligning with decreased use of the innervated legs. This decline may show functional loss, possibly resulting from nerve trauma during electrode implantation. As the surgical procedures improved and nerve damage was minimised, RMS values varied more by individual animal activity levels during measurement sessions. In several animals, RMS values increased after 1 to 3 days post-implantation, indicating of nerve recovery or swelling that enhances electrode-nerve contact.

Figures 5e-g provide detailed impedance and recording traces from locust 1 over the 30 days, with data for the other animals available in Figure SI8. All electrode impedances (Figure 5e) followed similar trends. Figure 5f illustrates recorded neuronal signals at three time points from both nerves. The blue trace represents the median signal recorded by the cuff electrodes. At the same time, the orange curve shows the bipolar signal between the electrodes at the edges of the cuff. Figure 5g highlights peaks identified in the signals, with contributing channels represented in different shades of blue. The bipolar traces illustrate how signals traveling in the opposite direction and generate distinct patterns. The direction can be afferent, which usually represents sensory signals, or efferent, which predominantly presents motor neuron signals. In the here displayed signals of locust 1, no bipolar signal was detected on the traces of day 28. Furthermore, the median peaks appeared broader. This broadening may result from other electric signals outside the cuff (e.g., EMG) or shorting between the electrode channels because of delamination. Observation of the animal showed it stopped using the leg around that time, indicating that the neuronal signals disappeared due to nerve damage. The exemplary traces of the other animals are shown in SI8.

All recordings, including those shown in Figure 5c-g, were performed while the locusts walked on an air-elevated polystyrene foam sphere (Figure 5h, Video SI10), allowing for natural movement and gait. We further analysed extended recordings from these sessions, to demonstrate our setup’s ability to capture behavioural differences in neural signals. The spikes in the median signal were identified using a negative threshold, identifying potential compound action potentials. At these time points, we also analysed the bipolar signal, which had distinct waveforms that were suitable for spike sorting. Principal component analysis (PCA) and k-means clustering were employed to categorise the shapes from a 4-minute-long recording, where the locust was both standing and walking. Figure 5i shows the first two principal components of the bipolar spikes, while Figure 5j displays the overlaid spike traces from each cluster. We chose 6 clusters for sorting. Additionally, we calculated the time shifts of the signals within each cluster between the first and last channel of the cuff (see Figure SI9 for details). The signal of cluster VI differs from others; it initially dips before reaching its positive peak. The propagation analysis within the cuff for this cluster revealed that these signals are predominantly afferent, suggesting a sensory origin. Compared to the others, cluster II primarily comprises signals with consistently low bipolar amplitude. These probably represent interfering signals originating from outside the cuff, which would explain their lack of propagation within the cuff, not resulting in a clear bipolar signal. However, time shift analysis also suggests the presence of some low-amplitude afferent signals within this cluster. The remaining clusters (I, III, IV, V) primarily represent efferent motor signals travelling towards the leg, distinguished by slightly varying waveforms and amplitudes.

Figure 5k presents two 5-second spectrograms of the neuronal signals’ power spectral density (PSD) for each leg, displayed during both standing and walking. These spectrograms, synchronised with video recordings, are also available in supplementary videos SI10 and SI11 for two animals. Spikes, previously detected and classified, are overlaid on the spectrogram corresponding to their respective leg. We plotted the absolute spike amplitude of the median signal. This approach was chosen because the amplitude of bipolar signals is a composite measure, influenced not only by spike amplitude but also by travelling velocity, direction, and waveform shape. Since these latter parameters are already characterised and represented by the classified clusters, plotting only the absolute amplitude offers a complementary perspective. Detected spikes are colour-coded according to their classified bipolar clusters. While the locust is standing, the spikes from clusters II, IV, and VI are predominant. The few spikes from cluster VI, along with some spikes from cluster II, likely represent the expected sensory signals. However, cluster IV also shows constant efferent activity, indicating ongoing motor signals despite the locust not moving. In contrast, walking reveals a broader variety of signal shapes, suggesting a more complex integration of sensory and motor activity.

The experiment shows that the implanted devices can reliably record neural activity and capture behavioural differences. It highlights the utility of self-foldable multi-electrode arrays for more precise and durable investigation compared to traditional hook electrodes used in small animal electrophysiology experiments.

### Transcutaneous port

To demonstrate the applicability of our USB-C connector neuroelectonic implant as a transcutaneous port, we conducted a proof-of-concept implantation in a rat. The sleeve surrounding the USB-C connector was designed to protrude through the skin, preventing water ingress and providing a grip when connecting to the external amplifier. Additional structures, such as suture loops or a mesh for skin fixation, were incorporated to anchor the implant securely in place. The device is shown in Figure 6a, featuring the USB-C connector on the right and suture loops at the bottom of the sleeve for attachment to the skin. It includes a 6 cm long feedline with an S-loop to route the wiring subcutaneously from the rat’s neck to the vagus nerve, providing strain relief and minimising tensile forces on the nerve. A 3 mm wide cuff was designed to fit the 250-300 µm diameter vagus nerve. The cuff featured eight PEDOT:PSS ring electrodes and a stainless-steel ground electrode placed subcutaneously and connected to the USB-C connector’s shield. Figure 6c shows the cuff positioned around the vagus nerve, while Figure 6d depicts the rat with the connector anchored on its back between the shoulder blades.

**Figure 6:**
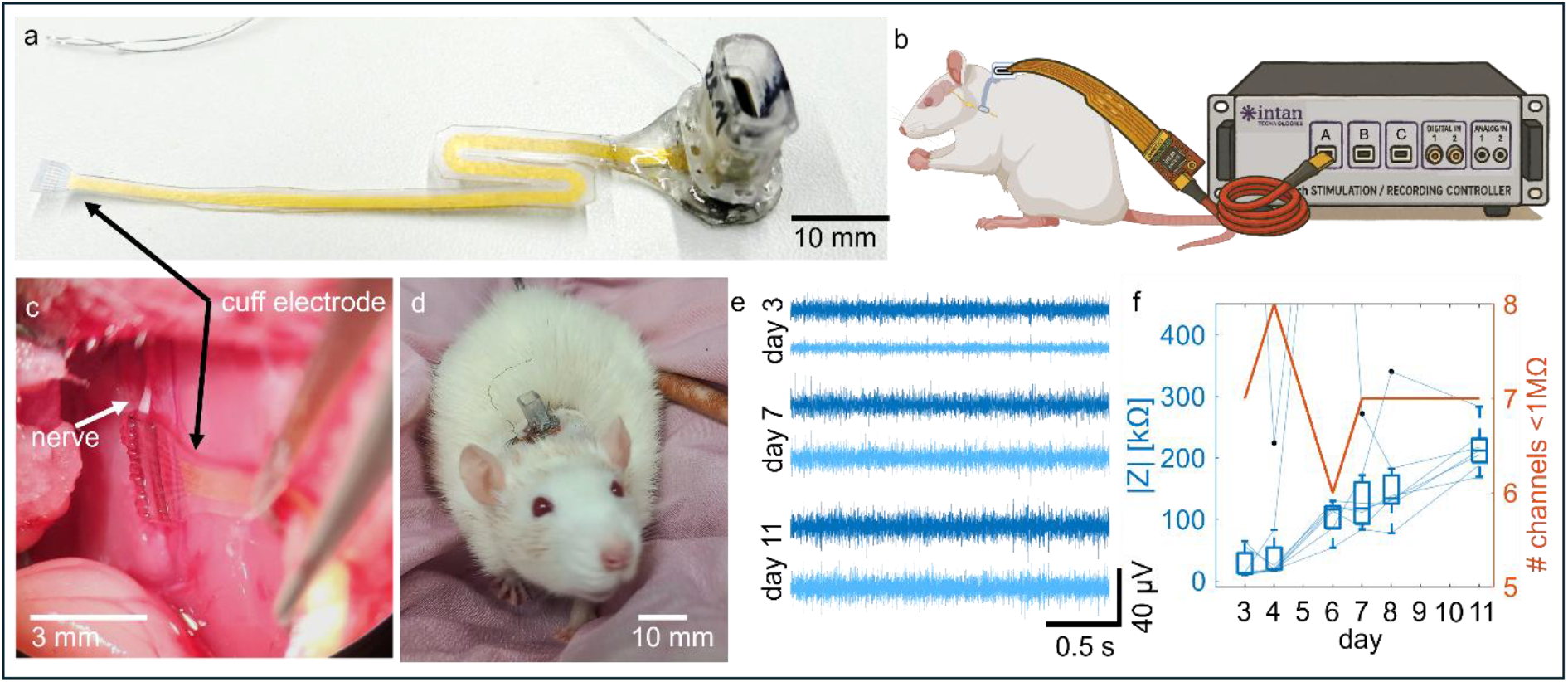
Implantation of self-folding electrodes with transcutaneous USB-C port for neural recordings on a rat vagus nerve. (a) Image of the self-folding electrode with an integrated USB-C connector designed for subcutaneous implantation in rats. The connector itself remains exposed through the skin, and S-shaped gold feedlines enhance mechanical decoupling between the connector and the nerve. (b) Schematic of the recording setup: the implanted electrode is connected via a flexible cable to the data acquisition headstage. (c) Image of the self-folding electrode placed around the vagus nerve of a rat. (d) Image of the implanted device in situ, with the USB-C port accessible through the skin. (e) Neural signal traces recorded from two cuff channels on days 3, 7, and 11 post-implantation. (f) Number of working channels and impedance measurements of electrode channels over 11 days post-implantation.

Chronic measurements (11 days total) commenced three days post-implantation to ensure healing and minimise the risk of wound opening at the connection site. The implant was connected using a flexible PCB to an Intan RHS32 headstage and an Intan RHS controller, as shown in Figure 6b. Figure 6e presents exemplary bandpass filtered (200-3000 Hz) signals recorded from two different electrodes on days 3, 7, and 11 after the implantation. As the animal was at rest during these recordings, the signals show the nerve’s characteristic spontaneous and tonic activity with the absence of discrete, high-amplitude spikes, which are typically only evoked by specific physiological stimuli. Figure 6f displays the channel impedances over time, together with the number of working channels (defined by showing an impedance <1 MΩ). The impedance increased more rapidly than in the locust experiments due to the pronounced foreign body response in mammals developing between the cuff electrodes and the stainless-steel ground wire. Nevertheless, most functional channels remained stable throughout the experiment. This example demonstrates that the devices are suitable for rodent experiments.

## Discussion

Recent years have seen significant progress in developing novel materials and fabrication techniques for neuroelectronic interfaces, with a focus on improving tissue compatibility by making them soft and flexible, lowering electrode impedance, and increasing channel count for enhanced spatial selectivity.^25,27–30,32–34^ However, in our opinion, a persistent bottleneck for the widespread adoption of these devices for long-term use by neuroscientists remains the lack of a robust, user-friendly method for connecting them to external data acquisition systems. This often results in delicate lead wires prone to failure, posing a significant barrier for long-term experiments, particularly in freely moving animals.^58^ Remarkably, most publications on neuroelectronic implants vaguely describe, if at all, their connection, and rarely show them.

Here, we demonstrate a fully integrated, ready-to-implant neural interface that addresses the critical challenge of establishing reliable electrical connections for chronic use with soft implants. We introduced a novel method for embedding rigid electronic components into the stretchable self-folding cuff electrodes for peripheral nerve interfacing. By utilising multi-material printing, we have mitigated electrical failure points between rigid components and the surrounding stretchable materials, creating a gradual stiffness transition that guides mechanical strain (see Figure 2e). This printing approach may also advance stretchable circuits beyond the neuroelectronic applications emphasised here.

We successfully integrated a wireless stimulation circuit and a standard USB-C connection port into the printed cuff electrodes. The monolithic integration of the USB-C port provides a “plug- and-play” solution, enabling reliable chronic nerve recordings. These devices not only simplify the surgical workflow due to the self-folding cuff but also preserve the integrity of the neural interface by decoupling the mechanical stress of plugging and unplugging from the nerve itself via the fixed connection port. While our approach is adaptable to various surface-mounted, vertically oriented connector types like the in neuroscience frequently used Omnetics Nano Strip series, we specifically chose USB-C for this demonstration due to its robust nature, widespread availability, and cost-effectiveness, making it an excellent showcase example for the stable transcutaneous port.

We believe that considering the whole system, including connectors, alongside material innovation, is essential for translating advanced neurotechnology from engineering labs to neuroscience research. The proposed solution for integrating a connector currently applies to printed self-folding cuff electrodes but might be adapted to other implant types. We hope it encourages researchers to address the entire system when developing new implants. In conclusion, we believe that our self-folding electrodes with a stable connection port will serve as a valuable tool for research in small animal models, particularly for studies involving freely moving animals, insect-machine interfaces, and insect cyborgs.^59–61^

## Materials and Methods

### Materials

The flexible 3D printing resins Luxaprint Flex and Medical Print Clear were purchased from Detax (Germany). The superabsorbent polymer (SAP) resin was mixed as previously described:^37,62^ Acrylic acid, 2-hydroxyethyl methacrylate (HEMA), sodium hydroxide (anhydrous; NaOH), poly(ethylene glycol) diacrylate (PEGDA, Mn 525), and 2,2,6,6-tetramethylpiperidine-1-oxyl (TEMPO), obtained from Sigma–Aldrich (USA) and the photoinitiator blend consisted of B(2,4,6-trimethylbenzoyl) phenylphosphine oxide/ethyl(2,4,6-trimethylbenzoyl) phenyl phosphinate (Omnirad 2100) and 2-isopropylthioxanthone (ITX) from IGM Resins (The Netherlands). Deionised (DI) water was generated by an Ultra Clear purification system (Evoqua Water Technologies, Germany), and 2-propanol (≥99.5%) was obtained from Carl Roth (Germany). For the PEDOT:PSS layer, ethylene glycol (EG), 4-dodecylbenzenesulfonic acid (DBSA), and (3-glycidyloxypropyl)trimethoxysilane (GOPS) were purchased from Sigma–Aldrich (USA), while the aqueous PEDOT:PSS solution (Clevios PH 1000) was obtained from Heraeus (Germany). Microscopy slides, used as carriers for the printing (76 × 52 × 1 mm^3^; Marienfeld) were purchased from VWR (USA). The locusts used in the study were purchased from a local pet shop.

### Fabrication

The devices were fabricated using frontal photopolymerization (FPP) of 3D printing resin mixtures, with additional steps based on a previously described fabrication method.^37^ First, the electrical components were inserted into a 3D-printed stabilising insert (Miicraft 50x, Miicraft Taiwan), typically made from the stiff medical print clear resin, or, if specifically stated, the flexible resin. This assembly was placed onto a roughened glass carrier and aligned using a USB camera. For part of the devices, e.g. the NFC circuit, a layer of PVA glue (local craft store) mixed 1:6 v:v with water was coated on the glass before placing the inlay, to avoid resin curing at the contact pads. Liquid flex resin (Luxaprint Flex, containing 0.4% w/w ITX as a UV absorber) was poured around the printed inlay across the rest of the print area. The structure was then exposed from below through the roughened glass using the Miicraft 50x 3D printer. After this first exposure step, excess uncured resin was removed using 2-propanol, and the glass slide was repositioned in the printer for the next layer. In this step, the SAP resin was selectively exposed in the hinge region to form swellable stripes aligned in the desired folding direction. Post-exposure, the unpolymerised SAP was washed off using DI water and 2-propanol. Before final UV curing, the printed structures were placed in an oven at 80 °C for 1 hour to relieve stresses. Final curing was performed in a nitrogen environment using an Otoflash G171 UV chamber (NK-Optic, Germany) with 2000 flashes, resulting in flexible substrates with integrated electrical components (Figure 4c). Following UV curing, the substrates were peeled off from the glass carriers, flipped, and transferred onto custom-made holders, ensuring the substrates lay flat. Care was taken to ensure that the printing resin did not obstruct the contacts; any residual resin on the contacts was carefully removed using scalpels under a stereomicroscope, or the protective PVA layer was washed off with DI water. The electrode tips were masked with polyimide tape to prevent metallization during subsequent processing. An 80 nm gold layer with a 10 nm titanium adhesion layer was deposited by magnetron sputtering (Moorfield nanoPVD, UK; 5 × 10^−3^ mbar argon atmosphere; 12 W Au, 40 W Ti). The sputtered metal areas were then treated with oxygen plasma to enhance surface hydrophilicity and promote uniform wetting. The PEDOT:PSS mixture containing 5 v/v% EG, 1 v/v% DBSA, and 2 v/v% GOPS was spin-coated onto the samples (Polo Spin 150i, SPS Europe B.V., Netherlands; 1500 rpm, 30 s). This coating procedure was repeated for three layers, each followed by a 1-minute drying at 110 °C on a hotplate. Before final curing, the electrodes were patterned using a nanosecond pulsed laser scanner (MD-U1000, Keyence, Japan; 500 mm/s scan speed, 18 µJ pulse energy, 80 kHz pulse frequency, 2 repetitions). The patterned devices were then fully cured on a hotplate at 110 °C for 1 hour to stabilise the PEDOT:PSS layer. For passivation, electrode sites were masked using polyimide tape, and a flex resin layer was spin-coated across the device surface (500 rpm, 30 s), sealing the conductive tracks and interconnects. Finally, the masking tape was removed from the electrode tips, and the final devices were cut along the grooves using scalpels. Implants and test samples with different geometries and designs were created using Inventor (2022, Autodesk, USA).

### Mechanical characterisation

For the mechanical test samples, the shape was designed in accordance with ISO 527 standards. Each sample featured a gauge length of 30 mm, a width of 5.8 mm and a thickness of 1 mm. Embedded within the samples were a red or blue LED (1206) alongside a 220 Ω resistor (0603), two 0 Ω resistors (0603) for conductance measurements, or no components for the blank samples. Mechanical testing was performed using a universal tester (Test 106, Test-GmbH, Germany) at a strain rate of 10% per minute, extending up to 20% elongation. Five-minute breaks were incorporated between cycles, with the testing environment maintained at 38°C, achieved using a regulatable halogen lamp. Resistance measurements were conducted against a 180 Ω resistor as voltage divider with an internal analog-to-digital converter. The setup included a custom tensile test clamp employing pogo pins for the electrical connections. The elastic modulus was calculated between 5% and 10% elongation and is reported as the mean and standard deviation of three samples, each with 11 cycles each. Finite element method (FEM) simulations were executed using COMSOL Multiphysics 5.4 (COMSOL, USA), assuming linear elastic material behaviour based on the previously measured values.

### NFC

The NFC circuit was designed around a commercially available SMD antenna and NFC chip. The antenna (ZC1003HF, Premo S.L., Spain) measured 10 x 10 mm^2^ and was tuned to the NFC operating frequency of 13.56 MHz using a 48 pF tuning capacitor (0603). An NTAG 5 chip (NTP53321, NXP Semiconductors, The Netherlands), optimised for energy harvesting applications, was used. The harvested output voltage was stabilised using a 2.2 µF and a 100 pF bypass capacitor (0603). An E4990A-120 Impedance Analyzer (Keysight, USA) with the 42941A Impedance Probe was used to measure the resonance frequency. The chip’s output voltage and the pulse width-modulated output were analysed with a digital storage oscilloscope (InfiniiVision DSOX2024A, Keysight, USA). In the centre of the devices, a thicker conductive gold layer of 100 nm was sputtered on top of the 80 nm layer (5 x 10^−3^ mbar argon, 12 W) to enhance conductivity. This area over the components was laser patterned with lower power (20% power, 500 mm/s scan speed, 80 kHz pulse frequency, 2 repetitions). A CLRC663 NFC evaluation board served as the reader, communicating with the integrated NTAG 5 chip via the NFC Cockpit software. A matched wire coil with a diameter of 45 mm and 6 turns, along with a 56 pF capacitor, was built to extend the effective communication range.

### USB-C Ports

For the embedded connectors, stainless steel (0.2 mm, 316L) wires were attached as ground electrodes to USB-C connectors (GCT USB4145-03-0170-C) and inserted into the 3D-printed inlay.

### Locust implantation

Adult male and female locusts (*Schistocerca gregaria*) were used for the in vivo experiments with the insect model. All experiments complied with the German laws for animal welfare (“Deutsches Tierschutzgesetz”). The surgical procedure used has been previously described.^37,63^ Before the surgery, the implants were fixed onto the animals using a light-curable band adhesive (Dental Technologies, USA), and locusts were anaesthetized by cooling them down to 4 °C for up to an hour. They were fixed onto a modelling clay bed, their ventral side up. By removing the cuticle around the metathorax and some of the underlying air sacs, the target nerve N5 exiting the metathoracic ganglion was accessed. In contrast to the previously described surgical procedure, the tracheae were not removed but slightly pushed away using glass capillaries to ensure they remained undamaged, which is critical for survival after the procedure. Self-folding electrodes were implanted around the nerves on both sides of the animals using fine tweezers. After successfully implanting the self-folding cuff electrodes on both sides, the cuticle was placed back on the cavity where the electrodes exit the body. The cavity was then sealed using melted beeswax (Fischbach Miller, Germany). The stainless-steel ground/reference electrode was placed in the animal’s abdomen. A custom-made polyimide flexible flat cable was used as the connection cable between the embedded USB-C connector and the data acquisition system’s headstage (RHS2116 INTAN Technologies, USA). During the data acquisition, the animals were placed on an air-elevated polystyrene foam ball, allowing recordings to be conducted on walking locusts. All recordings were done against the ground/reference electrode as a monopolar setup with a sampling rate of 30 kHz. Impedance measurements of the implanted channels were performed at 1 kHz with the same headstage, prior to signal acquisition. An LED connected to the digital output of the controller (128ch Stimulation/Recording Controller, INTAN Technologies, USA) was used as a trigger to synchronise the neural recordings with visual data acquired with a camera. Electrophysiology recordings were performed at various time points post-implantation over a 30-day period. The neural signals were processed using MATLAB (MATLAB 2021, MathWorks, USA). Signals from the working channels (|Z| < 1 MΩ) were bandpass filtered using a Butterworth filter between 200 and 4000 Hz. RMS values were calculated on 4-second data sections identified manually through synchronised video recordings. The data segments were selected from periods where the animal showed spontaneous movement, as determined by behavioural observation. PCA and k-means clustering approaches were used to derive 6 clusters of identified spikes. At the end of the implantation period, the implants were removed to check them. Therefore, the animals were anesthetised and fixed in the same manner prior to implantation. The beeswax seal and the cuticle covering the implant site were carefully removed. The self-folding electrodes were gently detached by slightly pulling them away from the interfaced nerve, releasing the nerve without rupture. Following explantation, the electrode tips were immersed in phosphate-buffered saline (PBS) for 1 hour and subsequently cleaned of any remaining tissue using fine tweezers.

### Rat implantation and electrophysiology

All experimental procedures were performed in accordance with the UK Animals (Scientific Procedures) Act 1986 and were approved by the animal welfare ethical review body at the University of Cambridge. These procedures were performed under a project license (PP5478947) by A. Güemes, with personal licence (I10076024) issued by the UK Home Office. An adult female healthy Lewis rat (Charles River Laboratories, UK), weighing 200–300 g, was used for rodent electrophysiology experiments. Animals were received at least one week prior to surgery for acclimation and were group-housed in individually ventilated cages with ad libitum access to food and water, maintained on a 12:12 h light-dark cycle (lights on at 07:00, off at 19:00). All procedures were conducted in accordance with the UK Animals (Scientific Procedures) Act 1986 under appropriate Home Office licenses and institutional ethical approval.

Prior to surgery, the rat was administered pre-operative analgesia (Carprofen, Carprieve® Injection, Norbrook Laboratories) along with a saline injection for fluid replacement. Anesthesia was induced at 5% and maintained at 2.5% using isoflurane inhalation. The animal was placed in a supine position, and a 15 mm longitudinal cervical midline incision was made to expose the left vagus nerve, following the procedure described by Güzel et al.^64^ The nerve was carefully separated from the carotid bundle via blunt dissection of surrounding perivascular tissue using finely curved forceps, with care taken to minimise manipulation and avoid nerve injury. Subsequently, the animal was repositioned dorsally, and a second incision was made between the scapulae down to the subcutaneous musculature. A subcutaneous tunnel was formed between the two incisions using blunt dissection. The implant wiring was threaded through the tunnel such that the microfabricated electrode portion rested at the cervical incision site, while the USB-C connector exited at the dorsal incision. The dorsal wound was sutured closed, and a custom-fabricated protective aluminium cap was secured over the connector to prevent damage from grooming or chewing. The electrode cuff was placed around the exposed vagus nerve at the cervical site, which was then sutured closed. Post-operative analgesia (Meloxicam, Metacam®, Boehringer Ingelheim) was administered twice daily for three days following surgery.

Recordings were performed on a maximum of five non-consecutive days, with a minimum of one rest day between series. Each recording session lasted at least 2 hours. On recording days, the rats were gently handled, the implant was connected via the USB-C port to a 32-channel headstage (Intan Technologies, Los Angeles, CA, USA), and the animals were placed back in their home cage to minimise stress. Neural signals were acquired at 20 kHz using the RHS2000 stimulation and recording system (Intan Technologies) and stored for offline analysis. We computed the average and standard deviation of the impedances for all working electrodes on each recording day and compared these metrics across different days. Channels that remained active across all recorded days were selected for further analysis. Signals from these selected channels were initially downsampled to 10 kHz and bandpass filtered (200 -3000 Hz) to isolate high-frequency neural components.^5,28,65,66^

## Supporting information

Video SI1 LED bend

Video SI3 LED bend modified

Video SI4 NFC stimulation

Video SI5 NFC LED

Video SI7 Free walking locust

Video SI10 Locust on Ball

Video SI11 Locust on Ball2

