## Supporting Information for "Integrating Electrical Components into a Printed Self-folding Cuff Electrode for Chronic Peripheral Nerve Interfaces"

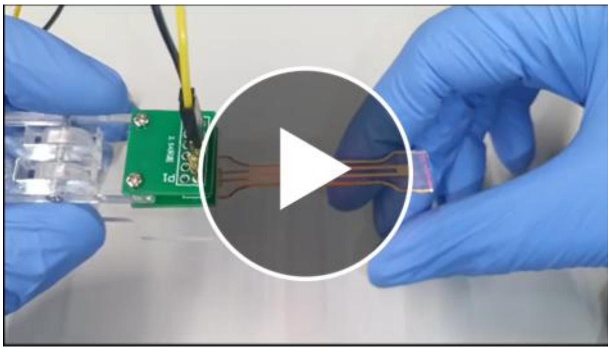

Video SI1: Demonstration of circuit failure in non-adapted devices due to bending. A printed test sample with an integrated LED is bent and the connection breaks irreversibly, where the inlay is made from the flexible and stretchable material.

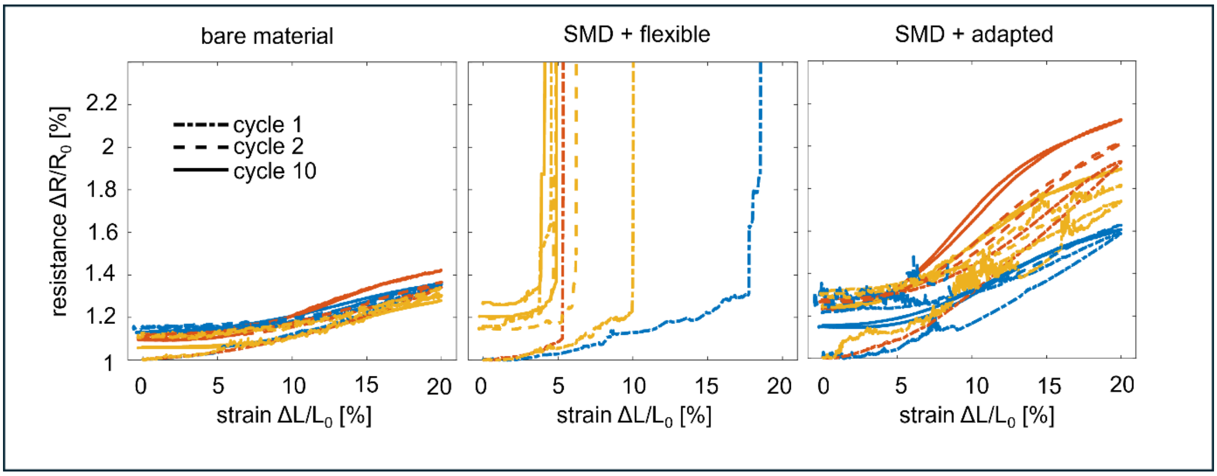


Figure SI2: Mechanical characteristics of test samples: Samples integrated with 0 Ω resistors during cyclic tensile testing up to 20% elongation. The relative resistance increase during the first, second, and tenth cycle is plotted over the strain.

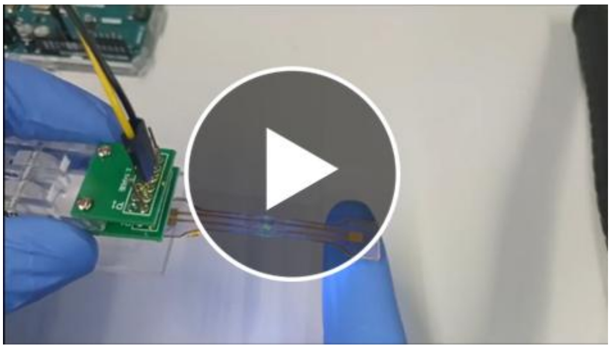

Video SI3: Bending of a stiffness adapted LED circuit. The test sample is printed with an inlay of stiff resin material, and the LED stays on during bending, showing intact connections.

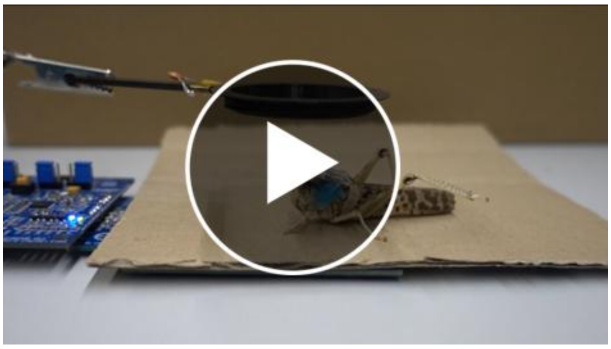

Video SI4: NFC stimulation: the N5 nerve of a locust is stimulated wirelessly using an NFC circuit for communication and power delivery. A distinct leg movement is visible during the stimulation pulse trains.

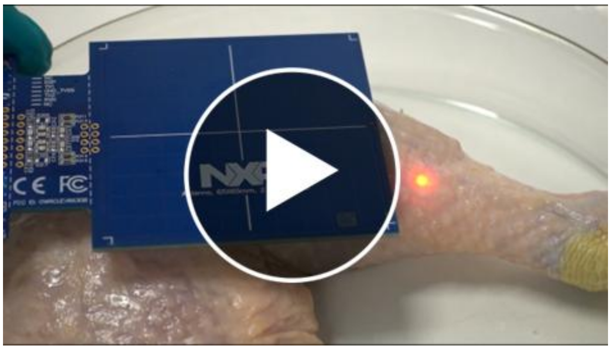

Video SI5: NFC LED circuit implanted: The stretchable NFC circuit with embedded LEDs is implanted under the skin of a chicken leg. The LEDs can be powered and controlled with the NFC reader board.

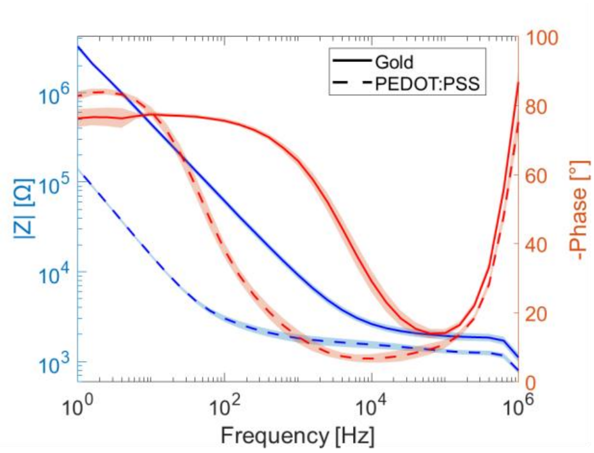


Figure SI6: Impedance spectroscopy of the self-folding electrodes with different electrode materials, gold and PEDOT:PSS, in PBS as electrolyte. Measurements were performed with a Palmsense4 potentiostat (Palmsens BV, Netherlands) vs. a Ag/AgCl reference electrode. A Pt coil electrode was used as a counter electrode.

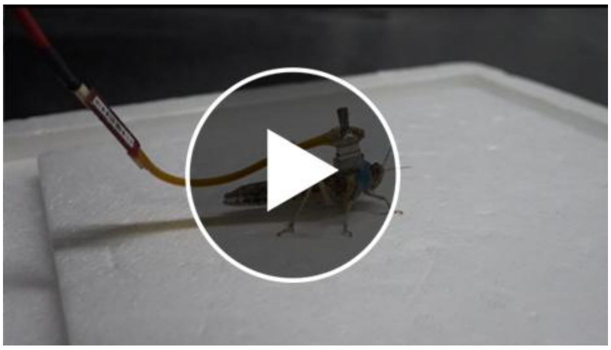

Video SI7: Free walking locust: Locust with the USB-C implant, connected to an Intan headstage via a custom FFC cable.

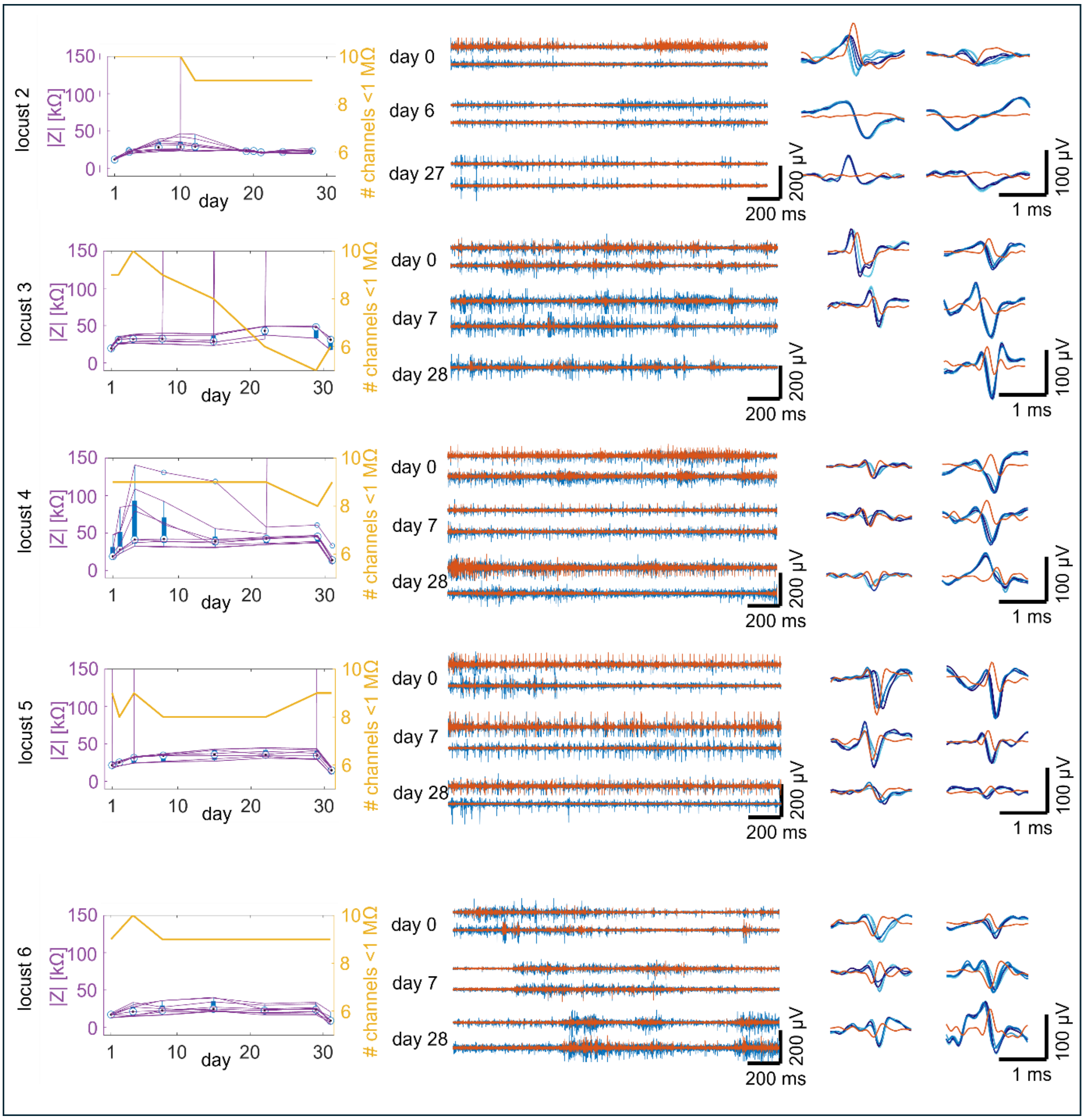


Figure SI8: Impedances and representative neuronal data at different days after implantation of additional locusts (supplementary to Figure 5).

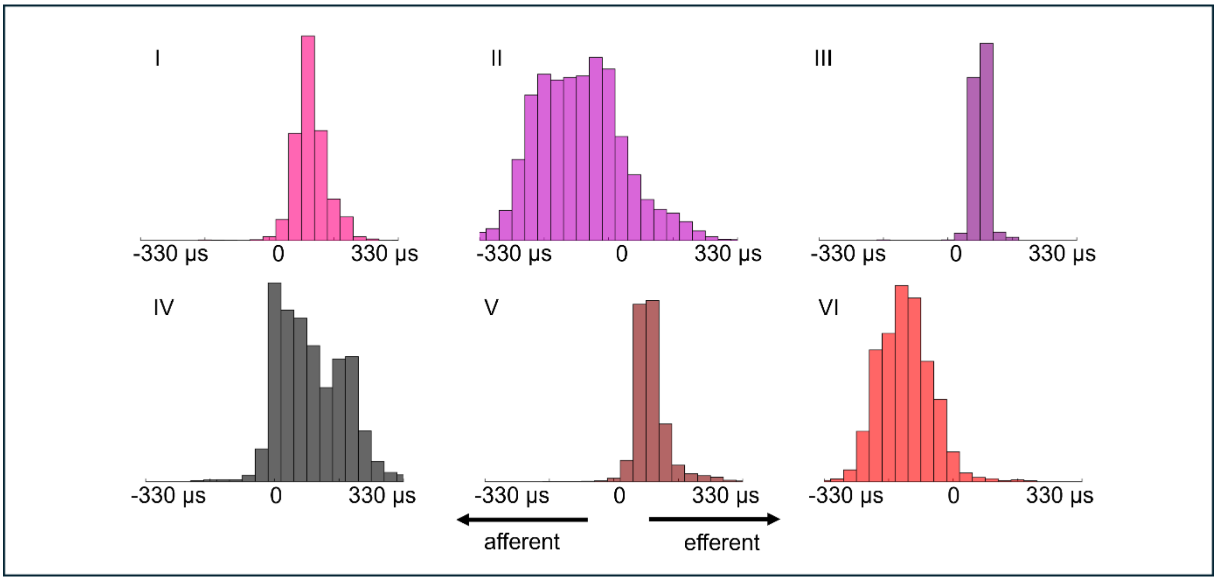


Figure SI9: Signal propagation of the identified spike clusters: Time lag of the signal between the most outer electrodes of the cuff (spaced 600 µm apart) revealing the propagation of the signals along the nerve. The shift was identified by the difference in the minima of the monopolar signals. Positive values indicate efferent activity, while negative values correspond to afferent signals.

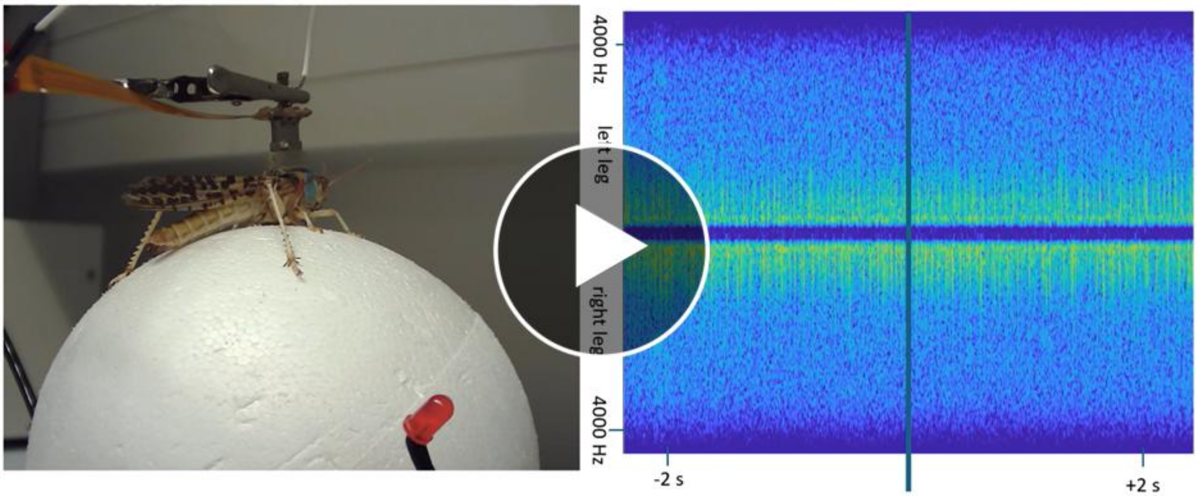


Video SI10: Locust on sphere, synchronised with the spectrograms of the power spectral density.

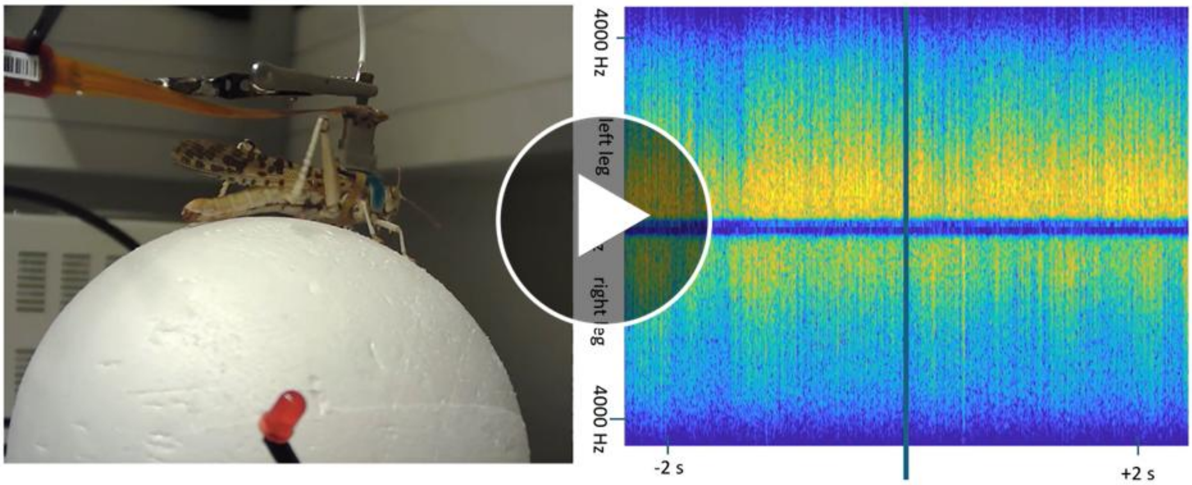


Video SI11: Locust on sphere, synchronised with the spectrograms of the power spectral density.
